# Microtubule Stabilization and Biomaterial Guidance Synergize to Enhance CST Regeneration and Motor Recovery After Chronic SCI

**DOI:** 10.64898/2026.02.05.703927

**Authors:** Usha Nekanti, Pooja S. Sakthivel, Rebecca A. Nishi, Andrea Anzalone, Courtney M. Dumont, Jeong Bin Lee, Skylar McDonald, Hui Song, André Obenaus, Paul David Gershon, Frank Bradke, Lonnie D. Shea, Brian J. Cummings, Aileen J. Anderson

**Affiliations:** Sue and Bill Gross Stem Cell Research Center, University of California, Irvine, CA, USA; Department of Anatomy and Neurobiology, University of California, Irvine, CA, USA; Department of Biomedical Engineering, University of Miami, Florida, USA; Basic Science Department, Loma Linda University School of Medicine, Loma Linda, CA, USA; Department of Orthopedics, Huazhong University of Science and Technology, Wuhan, China; Department of Biomedical Sciences, University of California Riverside, CA, USA; Department of Molecular Biology & Biochemistry, University of California, Irvine, CA, USA; Laboratory for Axon Growth and Regeneration, German Center for Neurodegenerative Diseases (DZNE), Bonn, Germany; Department of Biomedical Engineering, University of Michigan, MI, USA; Department of Physical Medicine and Rehabilitation, University of California, Irvine, CA, USA; Institute for Memory Impairments & Neurological Disorder, University of California Irvine, CA, USA

## Abstract

Spinal cord injury (SCI) results in loss of sensory and motor function below the level of damage, with chronic injuries presenting unique challenges for regenerative therapies. While multichannel biomaterial interventions have shown promise in promoting axonal regeneration, circuit restoration, and motor recovery in acute SCI, achieving similar outcomes in chronic injury models remains challenging due to a combination of intrinsic and extrinsic factors. These include the reduced capacity of the neuronal cell body to sustain a growth-activated state and the formation of a physical and chemical barrier at the injury site, preventing axonal growth. To address these challenges and promote motor recovery after chronic injury, we investigated the combinatorial effect of two regenerative approaches: 1) the implantation of poly (lactide-co-glycolide) (PLG) biomaterial bridge to guide axonal growth through the injury site, and 2) the delivery of Epothilone B (EpoB), a microtubule stabilizer that strengthens axons to promote regrowth. We used a transgenic mouse model that selectively expresses a red fluorescent protein variant (tdTomato) reporter throughout the corticospinal tract (CST) under control of the Crym promoter (Crym-tdTomato). We demonstrated that the combination of bridge implantation 60 days after surgical hemisection at C5 with EpoB improved locomotor function. At 12 weeks post-bridge implantation, immunohistology revealed axon regeneration in mice receiving implantation, but not EpoB or no-implant controls. The addition of EpoB significantly increased the volume of both total and CST axons regenerating through the biomaterial channels. Diffusion tensor magnetic resonance imaging (DTI) analysis identified enhanced fractional anisotropy (FA), axial diffusivity (AD), and mean diffusivity (MD) in the bridge region in the combination treatment group, consistent with new intact axons. Furthermore, EpoB enhanced the myelination of regenerated axons in the bridge. Finally, we investigated the proteomic profile of corticospinal neurons ipsilateral and contralateral to the SCI lesion and bridge, comparing the effect of EpoB treatment. Mass spectrometry-based analysis of laser-captured cells in this paradigm identified activation of a regeneration program by corticospinal neurons. These findings present a novel approach to enhance regenerative neural repair and locomotor recovery in chronic SCI.

## Introduction

Approximately 300,000 individuals in the USA live with SCI [1, 2]. The capacity of neuronal regeneration after an SCI is limited due to extrinsic cues deriving from cell death, axon loss, and demyelination [3, 4]. However, biomaterial interventions have proved promising for central nervous system (CNS) regeneration and locomotor recovery. Importantly, corticospinal tract (CST) axons are critical for the performance of forelimb functions in both rodents and humans, but are generally thought to exhibit the least potential of spinal tracts to regenerate after injury. Despite this intrinsically limited growth potential, we have shown that acute implantation of porous, biodegradable, multi-channel poly(lactide-co-glycolide) (PLG) bridges modulate the injury microenvironment and provide mechanical stability to drive neuronal regeneration, enabling CST regeneration. Indeed, CST axons regenerated and extended through the injury site, exhibiting synaptic connectivity with the triceps neuromuscular junction in association with motor recovery that was abolished after bridge transection [5, 6].

Critically, chronic SCI poses a different challenge due to both extrinsic and intrinsic barriers to regeneration. In addition to a molecularly hostile environment (extrinsic), neurons in the chronically injured cord are unable to sustain a growth-activated state (intrinsic) [7-11]. Although bridge implantation in the chronic period after SCI supports limited axon regeneration into the bridge, it does not improve motor function [12]. Here, we hypothesized that enhancing the intrinsic growth state of lesioned axons would drive regeneration in the chronic bridge implantation paradigm, enabling motor recovery.

Previous work has shown that microtubule stabilization via epothilone (Epo) administration can promote a more regenerative axonal state [13-15]. EpoB administration enables axonal growth through the injury site by inducing microtubule polymerization in the axon tip [16], a mechanism that drives axon growth during development [17]. Moreover, EpoB decreases fibrotic scarring and increases serotonergic fiber regeneration following a moderate contusion SCI [14, 17]. EpoB in combination with physical rehabilitation leads to enhanced gait recovery when compared to EpoB alone after SCI [14], highlighting its potential as a combinatorial treatment. We therefore sought to combine bridge implantation with EpoB administration following a chronic SCI to modulate both extrinsic and intrinsic barriers to CNS regeneration and enable motor recovery.

We implanted the bridge at 60 days post-hemisection SCI, followed by five systemic injections of EpoB starting on the day of implant and repeating every ten days **(Fig. 1)**. We used Crym-tdTomato reporter mice, which allow us to evaluate the CST axonal regeneration through bridge and investigate the regenerative profile of CST cell bodies in the motor cortex. We tested two different doses of EpoB: Low (0.75 mg/kg) and High (1.5 mg/kg). Our results showed that the combination of bridge implantation and EpoB delivery at both tested doses resulted in decreased paw placement errors on a horizontal ladder beam task at 12 weeks post-bridge implant compared to mice receiving either no treatment control, bridge, or EpoB treatment alone. Moreover, combinatorial bridge implantation and EpoB administration led to robust motor recovery and CNS regeneration, as quantified within the bridge by total axonal regeneration, CST axon regeneration, re-myelination of regenerated axons, and diffusion tensor magnetic resonance imaging (DTI) analysis. Lastly, laser capture microdissection of Crym-tdTomato+ corticospinal neurons from the motor cortex, followed by protein mass spectrometry and comparative proteomic analysis, demonstrated activation of a regenerative profile in these cells. Overall, our findings highlight the synergistic potential of combining bridge implantation with EpoB administration in chronic SCI, revealing a promising strategy that overcomes both extrinsic and intrinsic barriers to regeneration, leading to significant motor recovery and enhanced neural repair.

**Figure 1:**
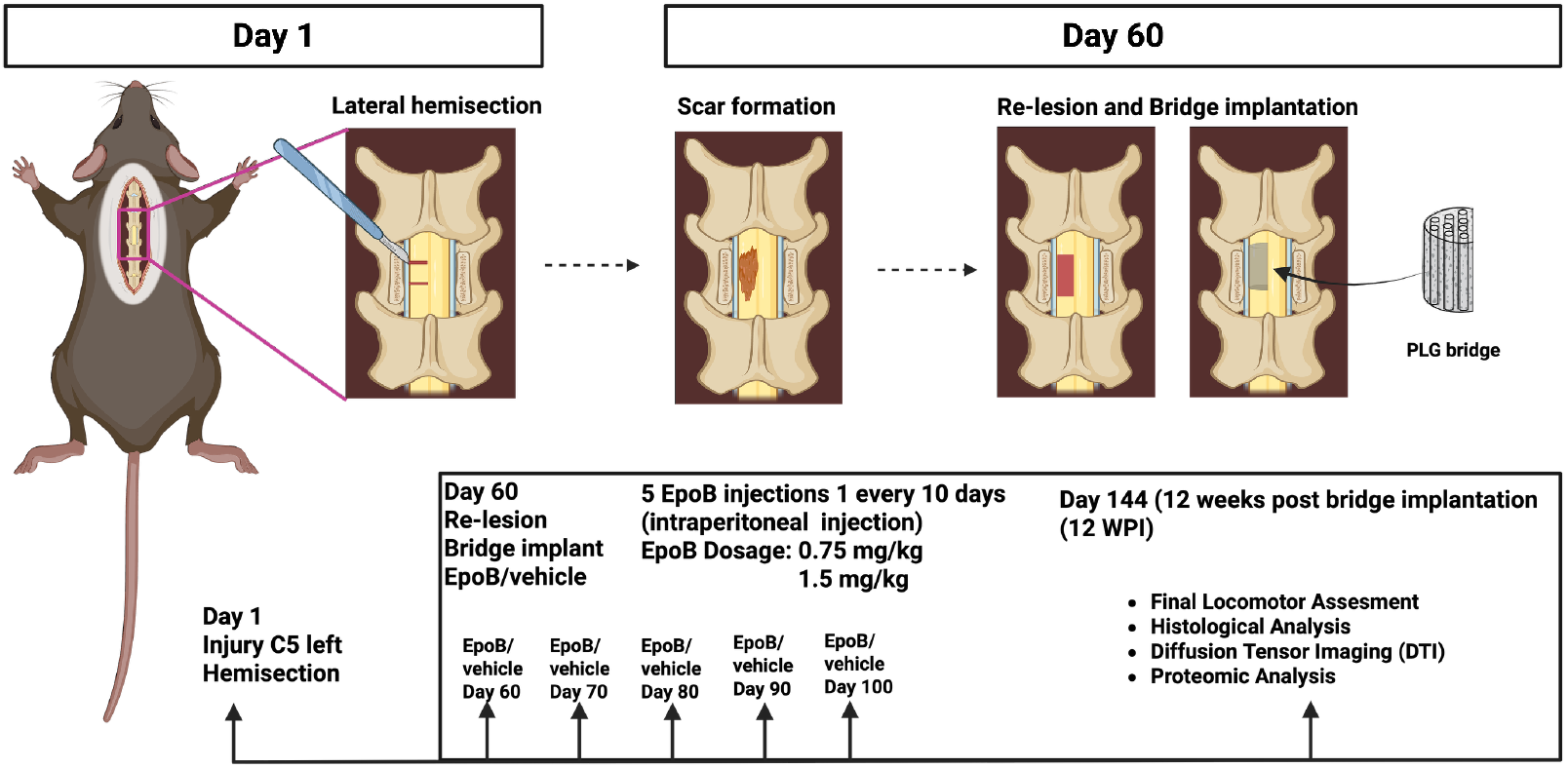
Experimental schematic. Timeline for the combinatorial approach of bridge implantation and Epothilone B (EpoB) administration in a chronic hemisection SCI mouse. Mice received a C5 double hemisection injury on the left, followed by re-laminectomy with scar tissue resection and bridge implantation at 60 DPI. Following bridge implantation, mice received an intraperitoneal injection of either 10% DMSO in saline (control), 0.75 mg/kg EpoB in 10% DMSO in saline (cohort 1), or 1.5 mg/kg EpoB in 10% DMSO in saline (cohort 2). Mice received a total of five control or EpoB injections, administered every ten days. Locomotor recovery was assessed using the horizontal ladder beam task at baseline (pre-injury), pre-implantation (60 DPI), and 4, 8, and 12 weeks post-implant. Mice were sacrificed at 12 weeks post-implantation and underwent either histological analysis to investigate axonal regeneration and myelination of regenerated axons within the lesion site ex vivo diffusion tensor imaging (DTI) to assess structural integrity and axonal organization within the lesion/bridge region, or proteomic analysis using laser-capture microdissection followed by mass spectrometry based proteomics to assess regeneration-mediated changes in corticospinal neurons ipsilateral and contralateral to the lesion. Animals treated with the 0.75 mg/kg dose (and corresponding vehicle controls) were used for immunohistochemical analyses and proteomics, whereas animals treated with the 1.5 mg/kg dose (and corresponding controls) were used for ex vivo DTI.

## Methods

### Ethics statements

Animal care, behavior acquisition, and data analysis were performed by investigators blinded to the experimental groups. All animal housing conditions, procedures, and animal care were approved by the UCI Institutional Animal Care and Use Committee (IACUC).

### Transgenic mouse derivation

The transgenic Crym^Cre^ mouse model was bred with mice harboring a targeted mutation with a loxP-flanked STOP cassette. Mu-Crystallin (Crym) is expressed in a diverse array of tissues, including layer V-VI of motor cortex in the brain; we have demonstrated that reporter expression is localized exclusively to CST axons in the spinal cord [6]. Crym^Cre^ mice will be crossed with Rosa Ai9 mice (JAX:007909), which carry a loxP-flanked STOP cassette at the Gt(ROSA)26Sor locus that blocks expression of a CAG promoter-driven red fluorescent protein variant tdTomato. Cre recombinase-mediated excision enables tdTomato expression (Crym^Cre: tdTomato^). Therefore, Crym^Cre: tdTomato^ exhibits red fluorescence specifically in CST axons in the spinal cord. All mice were group housed with 2-4 cage mates. Mice were randomly distributed into SCI control vs bridge groups, and into vehicle vs EpoB groups.

### Multichannel bridge fabrication

Multichannel PLG bridges were fabricated using the previously described gas foaming/particulate leaching method [12, 18]. Briefly, PLG microspheres were fused around salt and sacrificial sugar fibers to generate a porous scaffold containing nine longitudinal channels. Bridges were cut to a final size of 1.15 × 1.25 × 2.0 mm and stored in a desiccator until implantation.

### Spinal cord injury & animal care

Mouse C5 hemisection injury and postoperative care were performed as previously described [19]. Briefly, adult mice were anesthetized using 2% isoflurane for 5 minutes before surgery. All animals received C5 laminectomy and two unilateral incisions. 60 days post-injury (DPI), animals received a second surgery, scar resection, and removal of 1 mm hemisection spinal cord segment. Mice then received bridge implantations to fit the size of the cavity or control animals received no implant. Following the surgeries, 5-0 chromic gut was used to suture exposed muscle, and wound clips were used to close the skin. Post-operative care included mice being allowed to recover in cages on top of heating pads at 37°C overnight with clean Alpha-Dri bedding. Subcutaneous injections were administered postoperatively: Baytril (2.5 mg/kg) once a day for two weeks, lactated Ringer’s solution 1 ml once a day for five days, and buprenorphine (0.05 mg/kg) every 12h for three days. Bladder expression was performed twice daily until the end of the study.

### EpoB administration

Systemic Epothilone B (EpoB) dosing was performed as described previously [15]. EpoB (0.75 or 1.5 mg/kg; Sigma) was prepared in 10% DMSO in sterile saline; vehicle consisted of 10% DMSO in saline. Injections were delivered intraperitoneally immediately after the second surgery (re-laminectomy) and bridge implantation at 60 DPI. Four additional intraperitoneal doses were administered at 10-day intervals (for a total of five doses over 50 days from the initial administration). Animals treated with the 0.75 mg/kg dose (and corresponding vehicle controls) were used for immunohistochemical analyses and laser-capture microdissection followed by mass spectrometry-based proteomics, whereas animals treated with the 1.5 mg/kg dose (and corresponding controls) were used for ex vivo diffusion tensor imaging (DTI).

### Behavioral testing and locomotor recovery assessment

All experimenters were blinded when collecting and analyzing behavioral data. Prior to behavioral testing, animals were handled daily for two weeks to acclimate animals to human contact. Horizontal ladder beam and cylinder behavioral testing were performed to quantify motor recovery at baseline (pre-injury), pre-implantation, and 4, 8, and 12 weeks post-implantation. Animals were acclimated to the environment for five minutes before they were recorded walking across a horizontal ladder beam for three total trials. Video analysis quantified the percentage of left paw placement errors as previously described [20].

### Perfusion, tissue collection, and sectioning

At 12 weeks post-implantation, all animals received pentobarbital (100 mg/kg) before transcardial perfusion with 15 ml PBS and 100 ml 4% paraformaldehyde. Brain tissue and spinal cord segments (C3-C7) were dissected and cryoprotected overnight in 4% paraformaldehyde/20% sucrose. The tissue was flash frozen the next day using −60°C isopentane and stored at −80°C. 30 µm sections of spinal cord segments were made with a cryostat and Cryo-Jane tape transfer system (Leica Biosystems, Wetzlar, Germany). Slides were stored −20°C prior to immunohistochemistry.

### Immunohistochemistry

Slides were thawed at room temperature for 30 minutes before immersion in antigen retrieval solution (10 mM sodium citrate, 0.05% Tween 20, pH 6.0) within a 92-94°C water bath for 1h. Slides were cooled to room temperature and rinsed with water before staining. Slides were incubated with blocking buffer (1.5% donkey serum and 0.1% Triton X in PBS) for 1h and then primary antibody overnight. Primary antibodies used were: neurofilament-heavy chain (chicken, 1:500, Abcam) for axons and processes, tdTomato (rabbit, 1:500, Rockland) for CST, and MBP (mouse, 1:250, R&D Systems) for myelin basic protein. Following three five-minute washes (0.1% Triton-X 100 in PBS), tissue was incubated with fluorescent dye-conjugated secondary antibodies (Jackson ImmunoResearch, West Grove, PA) and nuclear dye Hoechst 33342 (1 µg/ml dilution; Invitrogen, Waltham, MA) for 2h. Slides were again washed three times and coverslips mounted with Fluoromount G (SouthernBiotech, Birmingham, AL).

### IMARIS histological analysis

Transverse spinal cord sections collected at 12 weeks post-implantation were used to quantify total regenerated and myelinated neurofilaments and CST (tdTomato) within the bridge. Triple immunohistochemistry was performed using NF-H (Neurofilament Heavy Chain, an axonal marker), MBP (Myelin Basic Protein, a pan-myelination marker), and tdTomato (CST). Quantitative analysis was performed as previously described [21]. Briefly, five to six random channels within the bridge were imaged using a Zeiss LSM 900 with Airyscan super-resolution microscopy at a 60X magnification. 3D surface volume rendering was performed using Imaris v9.6 (Oxford Instruments, Abingdon, United Kingdom). All three volumes of total neurofilament, total MBP-positive myelinated neurofilament, and total tdTomato reporter expression for CST were masked using the Surface feature. To exclude excess noise, a filter with a minimum voxel size of around 1000 was applied to each image. When masking the surface volumes. Next, the NF-H- and tdTomato-positive surface volume associated with myelin was masked using the object-to-object shortest-distance (0.4 µm) filter in Imaris.

### Diffusion tensor imaging (DTI)

Spinal cords were collected from mice perfused with 4% paraformaldehyde at 12 weeks following bridge implantation. DTI acquisition was performed on a 9.4T Bruker Avance MRI (Paravision 5.1) using a proton quadrature coil with the following parameters: TR/TE= 6500/35.6 ms, NEX=8, 25 slices at 0.5 mm thick, 1.5 × 1.5 cm field of view, 5 b0 and 30 directions at b=3078.18 s/mm2. Parametric DTI maps were generated using FMRIB’s Diffusion Toolbox from FMRIB’s Software Library (FSL), in which a diffusion tensor model was fit to each voxel. DSI Studio (https://dsi-studio.labsolver.org/) was used to generate deterministic fiber tracking using an angular threshold of 30 degrees and a step size of 0.03 mm. Fiber trajectories were smoothed by averaging the propagation direction with a percentage of the previous direction. Tracks with a length shorter than 0.5 or longer than 10 mm were discarded. A total of 500,000 seeds were placed to generate tracts and were seeded 3 mm above and below the lesion site. Streamline (tract) features were extracted and summarized in MS Excel.

### Nano Liquid chromatography mass spectrometry (nanoLC-MS/MS)

A cryostat was used to generate 14 µm brain sections on membrane slides (Zeiss 415190-9081-000). Approximately 150-250 CST neurons (tdTomato +) from layer 5 motor cortex (Bregma 2.4 - 1.4) were laser captured into adhesive cap tubes (Zeiss 415190-9201-000) using photoactivated localization microscopy (PALM) laser microdissection system (Zeiss Oberkochen, Germany). Tissue samples in the adhesive caps were solubilized in a minimum volume of 10 mM in tris(2-carboxyethyl)phosphine (TCEP), 0.1 M in triethylammonium bicarbonate (TEAB), 0.24% RapiGest then supplemented with LysC protease (50 ng/µl) then sealed into tubes for overnight incubation at 37°C. After supplementing with trypsin (25 ng/µl) for 6h incubation at 37°C, formic acid (FA) was added to 2% final concentration and peptides were desalted using stacked C18/SCX stage tips generated in house. Stage tip elutions were dried under vacuum and desalted peptides redissolved in 10 or 12 µl 0.1% FA in water. 30 micron ID x 360 micron OD x 75 cm columns were packed in-House with ReproSil-Pur C18-AQ beads (1.9 µm diameter; Dr. Maisch GmbH) and operated at 60^0^C. Using an EASY-nLC 1200 (ThermoFisher, Waltham, MA), columns were pre-equilibrated at 900 bar with 6 microL 0.1% FA in water (solvent A) then loaded at 900 bar with half of each desalted sample (5 or 6 microL) followed by a discontinuous gradient from 0 to 5% solvent B (93% CH3CN in 0.1% FA/water) over 5 min then to 20% solvent B over 325 min then to 35% solvent B over 30 min, at a flow rate of either 40 or 50 nL/min. Column eluate was electrosprayed into an LTQ Orbitrap Velos Pro mass spectrometer (ThermoFisher, Waltham, MA), acquiring FTMS precursor spectra from 380 - 1600 m/z (maximum ion injection time = 100 ms) at 60,000 resolution in positive ion profile mode followed by MS2 (FTMS, 7,500 resolution; maximum ion injection time = 300 ms) of the 15 most intense >+1 charged precursor ions with intensity > 1500 arbitrary units, fragmenting by HCD (30% NCE). Some acquisitions employed a reject mass list for MS2 with 646 distinct keratin m/z values at 10 ppm mass width. After two fragmentations within a 30 sec window, ions were dynamically excluded, via a 500-entry list, for 40 sec or until rolling off the list by surpassing the maximum number of entries. Early expiration from the list was enabled for a S/N threshold of 2.0.

### Analysis of nanoLC-MS/MS data

MS raw files were imported to MaxQuant [22]. The Andromeda search engine was set with a maximum of 2 missed trypsin sites, minimum peptide length = 7 residues and default peptide modifications. Quantitation method was label-free, normalization type was “Classic”. Search databases were UniProt Mouse (forward and reverse), the default “common contaminants” database, a second contaminants database to include recombinant LysC and a custom database for TdTomato. The “match between runs” option was used with match time and alignment windows of 2 and 85 minutes, respectively. Results were thresholded at 2% and 4% peptide and protein level FDR, respectively. Successful protein quantitation required at least two peptide identifications (razor + unique) per protein. “Protein groups.txt” output from MaxQuant was uploaded to “Perseus” [23], whose matrix was filtered to exclude common contaminants, hits to the reverse database and proteins identified only via modified peptides. Sample groupings were defined as follows: contralateral EpoB, contralateral Bridge + EpoB, ipsilateral EpoB, and ipsilateral Bridge + EpoB. Quant values were converted to log2 then filtered for the successful quantitation of >2 valid values in total. Missing values were imputed based on a normal intensity distribution (width: 0.3; downshift: 1.8) then, as a normalization step, median intensity was subtracted from each value in the “intensity data” column. Log2(per-protein-mean-quant-value-within-sample-grouping) were compared on a pairwise basis by volcano plot.

Differentially expressed proteins were identified from protein hits using a cut-off of p < 0.05 by Student’s T-test. Gene ontology by over representation analysis was run on significant protein hits to determine cellular compartments that were altered in each comparison of interest. Protein hits were also evaluated to a previously published dataset (GSE126957) to establish relevance of proteins to enhanced neuronal plasticity.

### Statistical analysis and exclusion criteria

Statistical analyses were performed using GraphPad Prism version 9.2.0. Mice were randomized across groups to ensure an unbiased distribution by age, sex, and weight by an investigator not directly involved in the study. The sample size (n), statistical tests conducted, and statistical significance are indicated in the respective figure legends. All data were expressed as mean ± standard error of the mean (SEM). Data collection and analysis were performed by investigators blinded to experimental group assignments. Unpaired Student’s t-tests were applied when comparing two experimental groups. ANOVA analysis was applied when comparing three or more groups. No mice were excluded from ladder beam analysis, immunohistochemical analysis, or proteomic analysis **(Fig. 2, 3, 4, and 6)**. For DTI analysis, one mouse from the Bridge + EpoB group was excluded due to an unclear lesion boundary **(Fig. 5)**.

**Figure 2:**
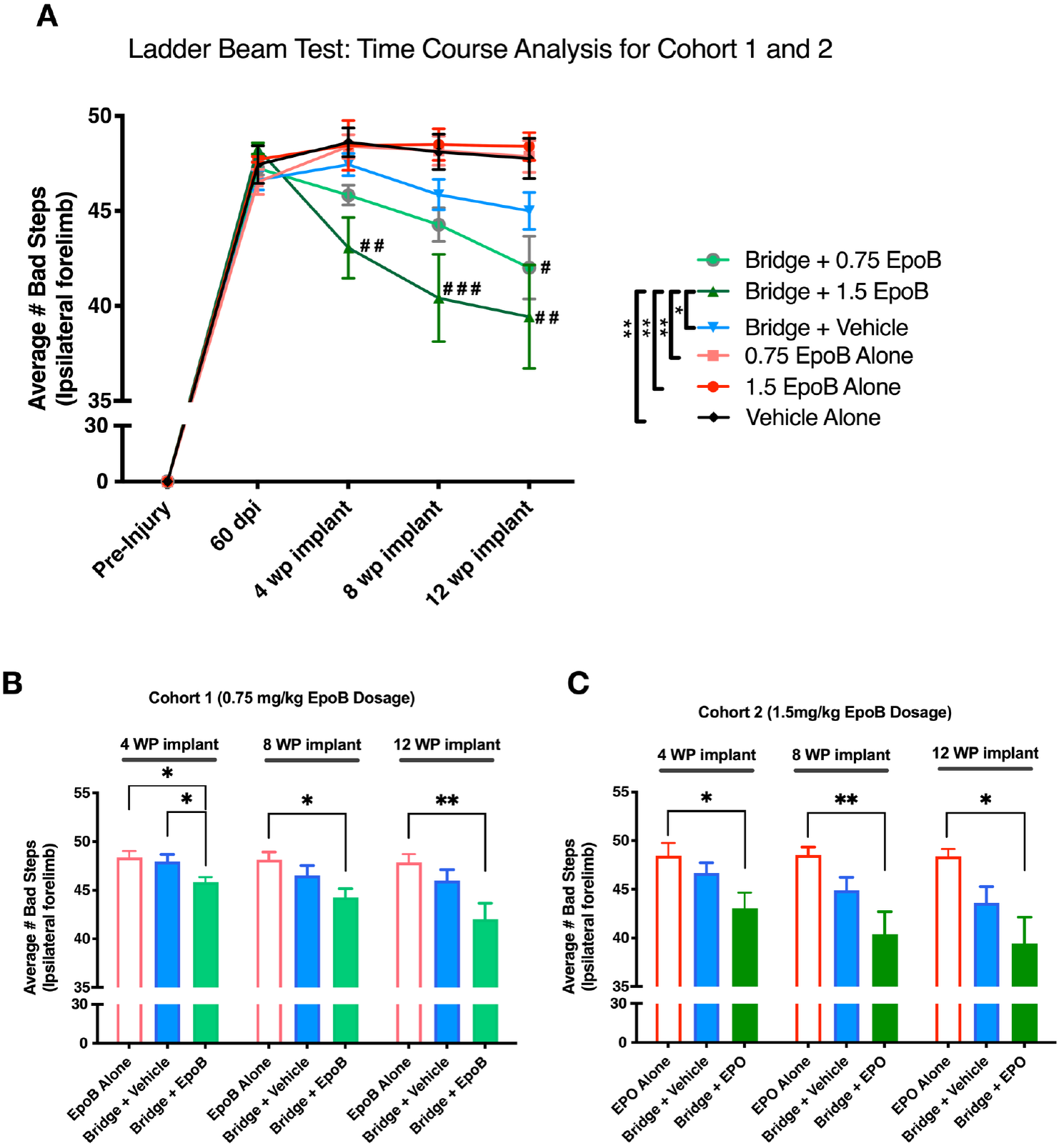
Combined Bridge implantation and Epothilone B treatment synergistically enhance locomotor recovery after chronic SCI in a dose-dependent manner. Locomotor function was assessed by quantification of ipsilateral forepaw placement errors per 50 rungs traversed on a horizontal ladder beam. **(A)** Time course analysis depicting locomotor performance at pre-injury baseline, pre-implantation, and 4, 8, and 12 weeks post-implantation, comparing control vs treatment groups from both cohort 1 and 2. * indicates Two-Way ANOVA followed by Tukey’s multiple comparison test (*P≤0.05, **P≤0.01). # indicates One-Way ANOVA followed by Tukey’s multiple comparison test performed at individual time points (#P≤0.05, ##P≤0.01, and ###p≤0.001). **(B-C)** Comparison of ipsilateral forepaw placement errors in 0.75 mg/kg EpoB dose (Cohort 1; **B**) and 1.5 mg/kg EpoB dose (Cohort 2; **C**) at 4, 8, and 12 weeks post-implantation (12, 16, and 20 weeks post-injury), showing significant improvement in the Bridge + EpoB group. **\* (B-C)** indicates One-Way ANOVA followed by a Tukey multiple comparisons test at each time point. All graphs represent Mean ± SEM; n = 10-12 mice/group in cohort 1 **(B)** and n = 6-7 in cohort 2 **(C)**.

**Figure 3:**
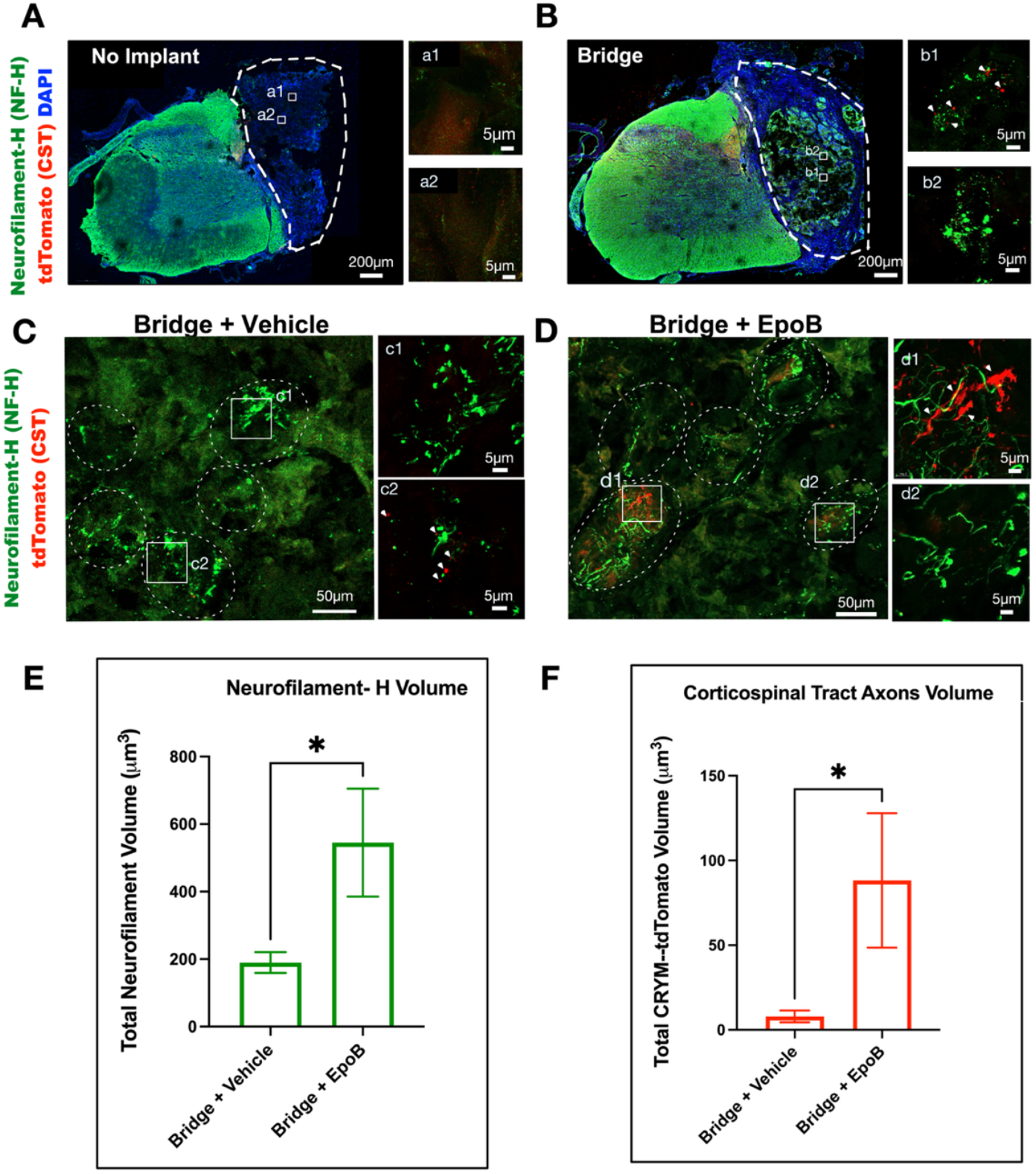
Bridge-mediated axonal regeneration is significantly enhanced by EpoB treatment in chronic SCI at 12 WP implantation. Representative immunofluorescence images and quantification of axonal regeneration within the lesion site. (A-D) Neurofilament-H (NF-H, green) to label total axons and CYRM-tdTomato (red) to identify corticospinal tract (CST) axons, with DAPI (blue) as a nuclear counterstain in transverse spinal cord sections. (A-B) White dashed lines outline the lesion and tissue interface. (A) Transverse spinal cord sections from no-implant control mouse, (a1-a2) higher-magnification views of the lesion site. (B) Transverse spinal cord section from a mouse with bridge implantation, (b1-b3) higher-magnification views of bridge channels containing NF-H and CST-labeled axons. White arrowheads in (b1) indicate CST axons. (C-D) Representative images showing axonal growth, including CST axons, in the transverse section of the bridge and within the bridge guidance channels (circled in white dashed lines, denoting the channels in the transverse bridge section). (C) Transverse spinal cord sections of mouse with bridge implant from Bridge + Vehicle group, (c1-c2) higher-magnification views of channel regions containing NF-H and CST-labeled axons. White arrowheads in (c2) indicate CST axons. (D) Transverse spinal cord sections of mouse with bridge implant from Bridge + EpoB group, (d1-d2) higher-magnification views of channel regions containing NF-H and CST-labeled axons. White arrowheads in (d1) indicate CST axons. (E) Quantification of total NF-H positive (NF-H+) axon volume within the bridge channels, comparing Bridge + Vehicle and Bridge + EpoB treated groups. (F) Quantification of tdTomato-positive CST axon volume within the bridge channels in Bridge + Vehicle and Bridge + EpoB-treated groups. Data are presented as Mean ± SEM; N=5 mice/group. **\*** Indicates unpaired t-tests (*p≤0.05). Scale bars are as indicated.

**Figure 4:**
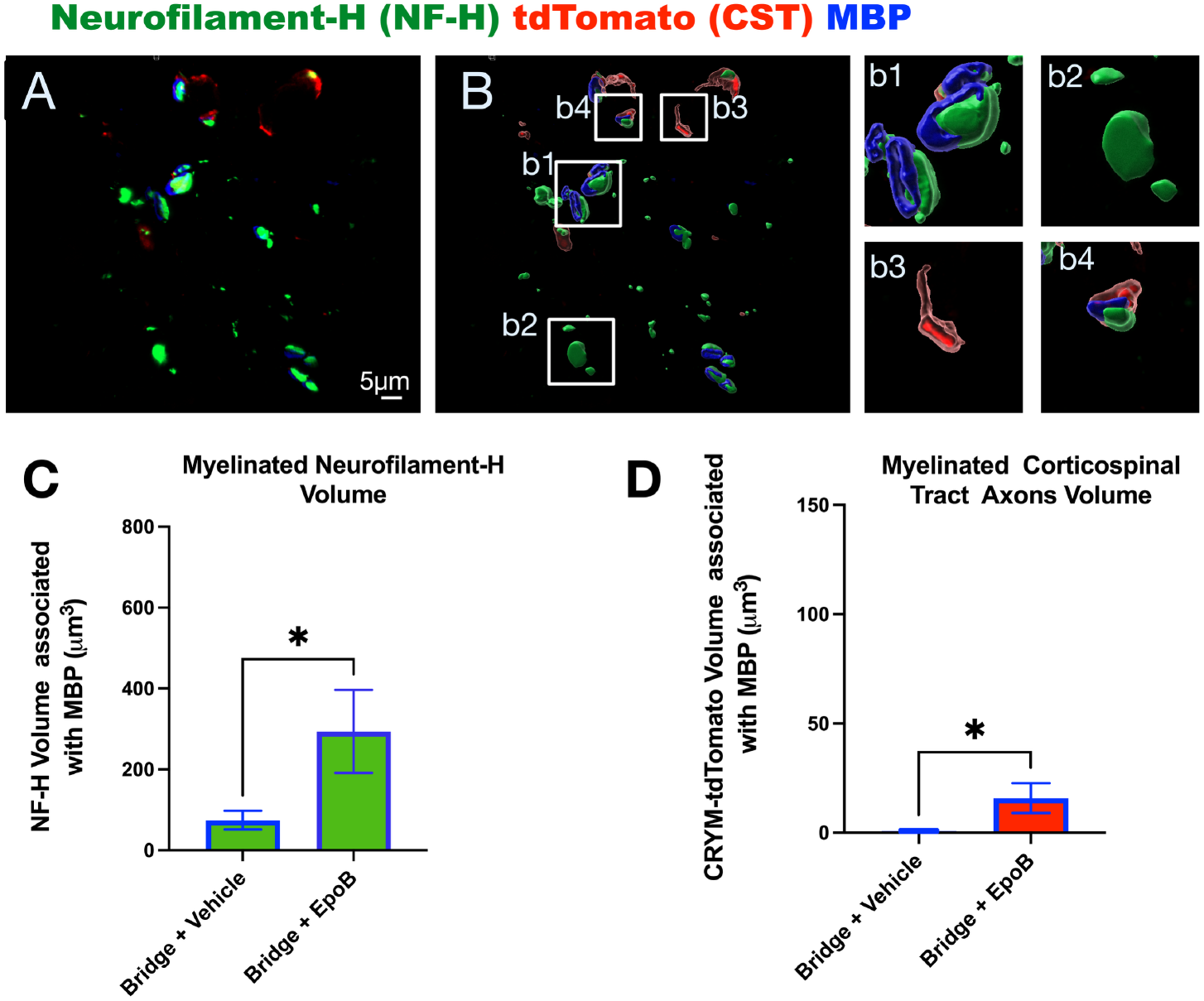
Combination of bridge and EpoB administration promotes myelination of regenerated axon. Immunofluorescence staining and quantification of myelinated axons within the bridge channels at 12 weeks post-implantation. (A) Representative image showing triple immunostaining for neurofilament-H (NF-H, green), CST axons (tdTomato, red), and myelin basic protein (MBP, blue) within the bridge channels implanted at the spinal cord lesion site. (B) Imaris 3D surface volume rendering of regenerated axons. Inset images provide higher magnification examples: (b1) myelinated NF-H+ axon, (b2) unmyelinated NF-H+ axon, (b3) unmyelinated tdTomato + CST axon, and (b4) myelinated NF-H+ tdTomato+ (CST) axon. Scale bars are as indicated. (C) Quantification of myelinated NF-H+ axon volume (NF-H volume associated with MBP) in Bridge + Vehicle and Bridge + EpoB groups. (D) Quantification of myelinated tdTomato + CST axon volume (Crym-tdTomato volume associated with MBP) in Bridge + Vehicle and Bridge + EpoB groups. Data are presented as Mean ± SEM; N=5 mice/group. **\*** indicates unpaired t-tests (*p≤0.05).

**Figure 5:**
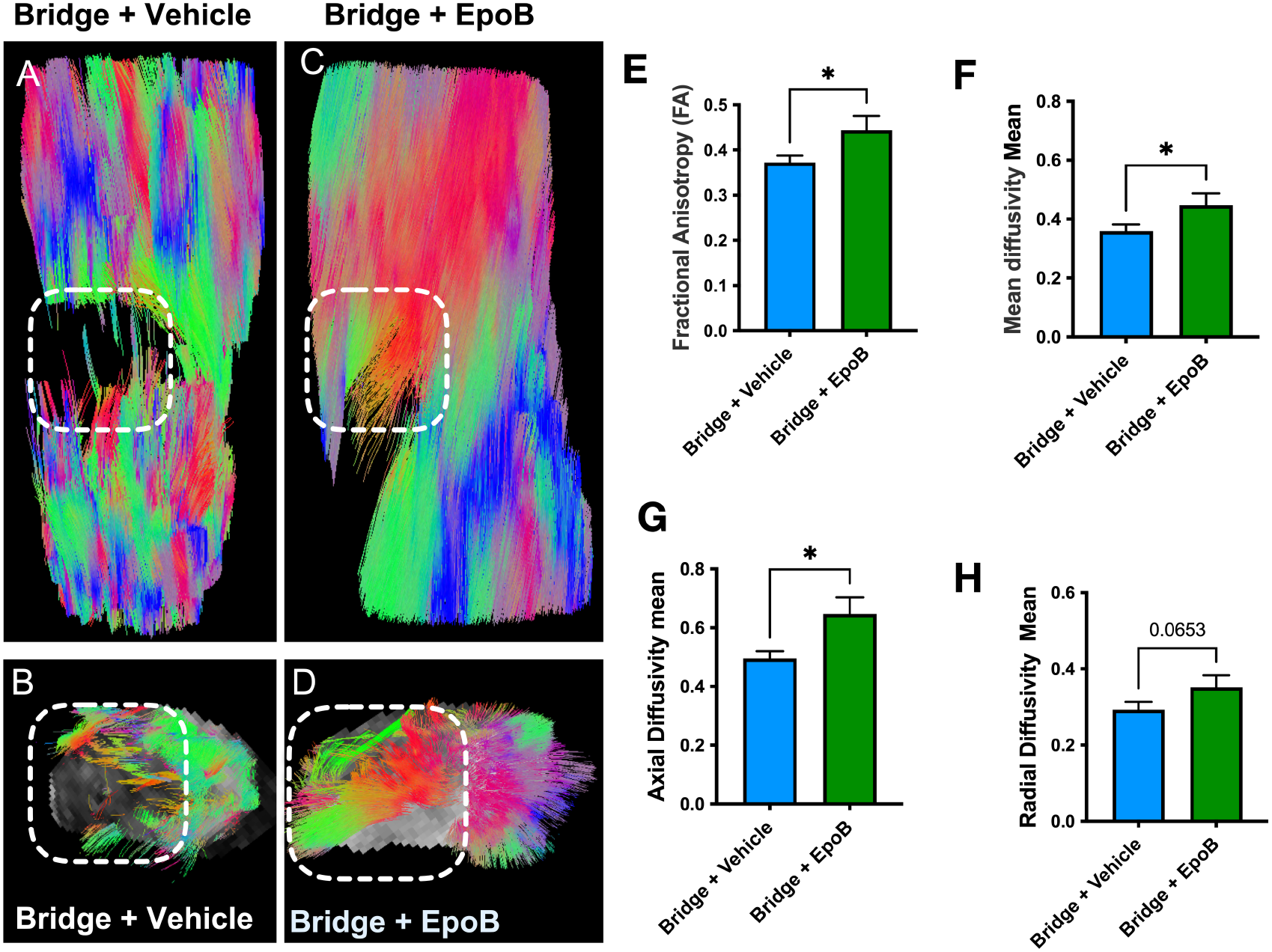
DTI analysis reveals improved axonal integrity with Bridge and EpoB treatment after SCI. Diffusion Tensor Imaging (DTI) analysis of mouse spinal cords at 12 weeks post-bridge implant and EpoB treatment (1.5 mg/kg dosage). (A-D) Representative DTI tractography reconstructions. Dashed white outlines denote the biomaterial bridge region. (A-B) Longitudinal (A) and transverse (B) views of a spinal cord treated with the bridge and vehicle control, showing relatively sparse tracts within the bridge. (C-D) Longitudinal (C) and transverse (D) views of a spinal cord treated with the bridge and EpoB, demonstrating a denser and more organized tract within the bridge area compared to the Bridge + Vehicle control. (E) Fractional anisotropy (FA) was significantly increased in the Bridge + EpoB group compared to Bridge + Vehicle. (F-H) Diffusivity metrics quantified within the lesion/bridge region: (F) mean diffusivity (MD), (G) axial diffusivity (AD), and (H) radial diffusivity (RD). The Bridge + EpoB group showed significantly higher MD and AD, while RD showed a trend toward increase (p=0.0653) compared to Bridge + Vehicle. N=8 mice in the Bridge+Vehicle group and N=6 mice in the Bridge + EpoB group. Data are presented as Mean ± SEM. Asterisks indicate unpaired Student’s t-tests (*p ≤ 0.05).

**Figure 6.**
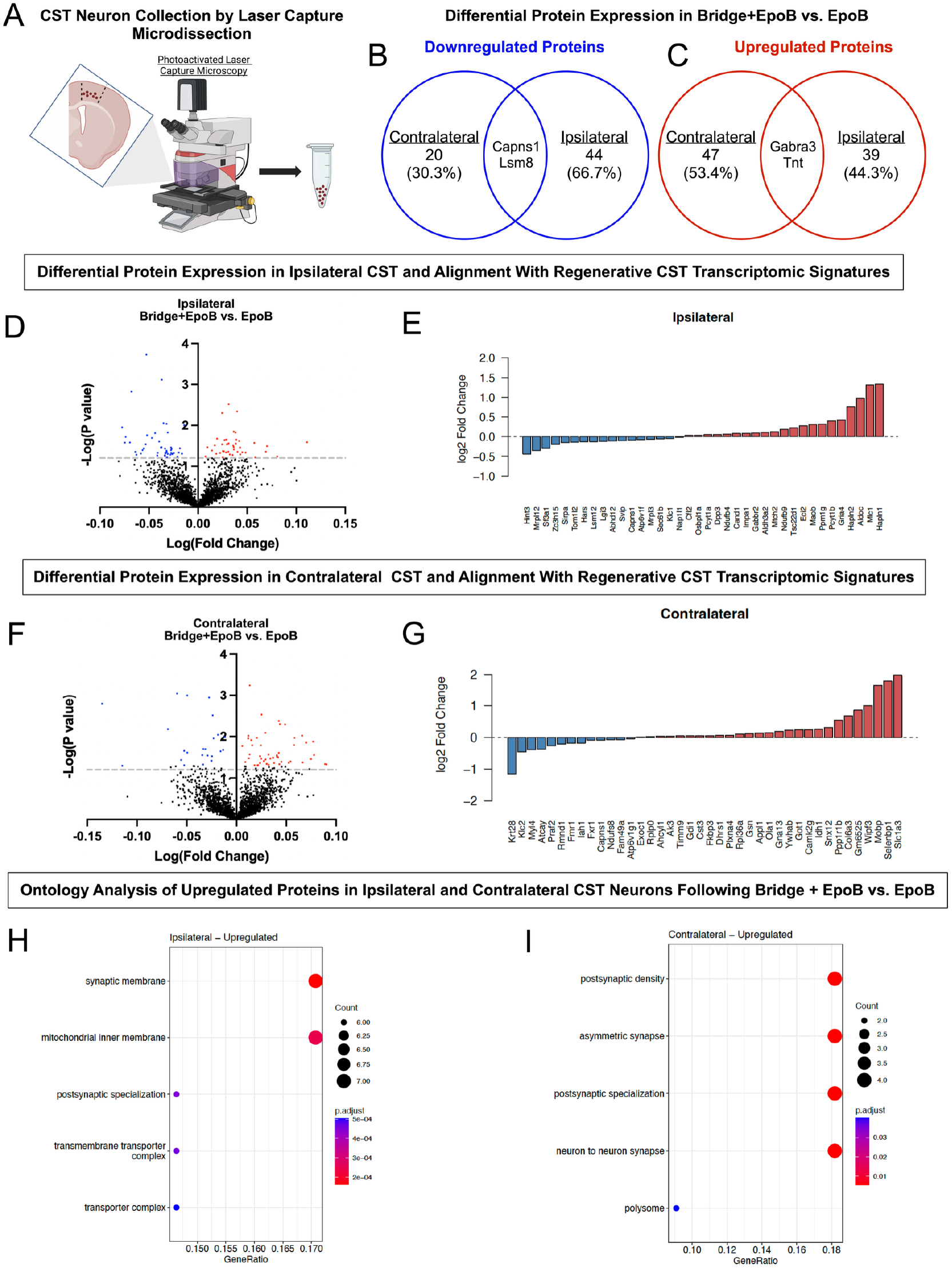
Proteomic analysis reveals increased pro-regenerative phenotype in CST neurons following bridge and EpoB administration, when compared to EpoB alone. (A) Coronal brain section of CrymCre-tdTomato mice highlighting layer V CST neurons that were collected by laser capture microscopy before analysis by lnanoLC-MS/MS. (B-C) Distinct protein signatures were identified in both the ipsilateral and contralateral cortices with minimal overlap in the significantly upregulated (B) and downregulated (C) proteins. (D-G) Volcano plot demonstrating the significantly upregulated (red) and downregulated (blue) proteins in Bridge + EpoB versus EpoB alone in the ipsilateral (D) and contralateral (F) hemispheres. Many differentially expressed proteins exhibited similar upregulation or downregulation at the transcriptome level in both the ipsilateral (E) and contralateral (G) hemispheres, as confirmed by comparison to a pre-existing dataset of pro-regenerative CST neurons from Poplawski et al. (2020). (H-I) Ontology analysis of the significantly upregulated proteins in our data reveals that PLG + EpoB, compared to EpoB alone, results in an upregulation of synaptic proteins in the ipsilateral (H) and contralateral (I) hemispheres. N=3 mice in the EpoB alone group and N=4 mice in the Bridge + EpoB group.

## Results

### Bridge implantation at 60 days post-SCI in combination with EpoB administration improves locomotor recovery in a synergistic manner

As described previously for acute bridge implantation in this model, mice receiving chronic bridge implantation were assessed for locomotor recovery using the horizontal ladder beam by scoring paw placement errors. No significant improvement in ipsilateral paw recovery was observed in either the no-treatment control or single-treatment groups. However, by 12 weeks post-implant (20 weeks post-injury) mice receiving either Bridge + 0.75 mg/kg EpoB, or Bridge + 1.5 mg/kg EpoB, performed significantly better than controls (Fig. 2A). Two-Way ANOVA followed by Tukey multiple comparison test with treatment groups and time point of assessment as a variable demonstrated that only Bridge + 1.5 EpoB showed significant improvement compared to all other treatment groups **(Fig. 2A)**. Breaking down this analysis to evaluate low and high dose EpoB treatment separately identified that both dose cohorts showed synergistic locomotor recovery, which was evident as early as 4 weeks post-implant **(Fig. 2B-C)**. Overall, treatment with 1.5 mg/kg EpoB exhibited a trend for more significant recovery vs. control groups than 0.75 mg/kg EpoB **(Fig. 2A-C)**. Together, these data demonstrate a distinct benefit of combinatorial treatment pairing bridge implantation and EpoB treatment on recovery of function in this chronic injury paradigm.

### Bridge implantation in chronic SCI supports CST regeneration and is augmented by EpoB

To investigate whether recovery of function was associated with axonal regeneration, immunohistological analysis was performed at 12 weeks post-bridge implantation using the 0.75 mg/kg EpoB cohort. Total axonal regeneration was assessed by immunofluorescent labeling for neurofilament-heavy (NF-H), while CST axons were identified using the Crym-tdTomato reporter, which selectively labels CST fibers in the spinal cord. Mu-Crystallin (Crym) protein is specifically expressed in CST axons within the spinal cord. Cre recombinase excises a floxed STOP cassette, resulting in tdTomato expression in these CST axons. Mice without bridge implantation did not exhibit NF-H positive (NF-H+) axons or Crym-tdTomato+ CST axons within the lesion site at 12 weeks post-implantation **(Fig. 3A, a1-2)**. In contrast, mice receiving delayed bridge implantation at 60 dpi displayed both NF-H+ axons (green) and Crym-tdTomato+ CST fibers (red), which were selectively localized within the bridge channels **(Fig. 3B, b1-2; Fig. 3C, c1-2;)**. Notably, this regenerative response was markedly enhanced in animals treated with both Bridge + EpoB **(Fig. 3D, d1-2)**.

Because axonal regeneration was observed exclusively in bridge-implanted animals, quantitative analyses were limited to comparisons between Bridge + Vehicle and Bridge + 0.75 mg/kg EpoB groups. These data confirmed not only that EpoB significantly enhances axonal regeneration; it revealed that EpoB treatment induces axonal regeneration even in the chronic phases of SCI, including the CST tract, which is known to be a notoriously poor regenerator. The volume of both NF-H+ **(Fig. 3E)** and Crym-tdTomato positive CST **(Fig. 3F)** axons within the bridge was significantly greater in the combination group compared to the Bridge + Vehicle group. These findings suggest that while the bridge itself provides a permissive scaffold for axonal growth, EpoB treatment further promotes the regenerative capacity of both total and CST axons through the injury site in the chronic injury environment.

### The volume of myelinated regenerated axons in bridge channels increased with EpoB treatment

Remyelination is crucial for functional circuit reintegration, particularly in long-tract pathways such as the CST, where majority of the axons are myelinated, as it restores conduction velocity and provides metabolic support essential for sustained function [24-26]. Based on the increased axonal regeneration observed with combination treatment, we next assessed the myelination status of regenerated axons using triple immunostaining for NF-H (total axons), tdTomato (CST axons), and MBP (myelination status) **(Fig. 4A)**. This analysis was again conducted within the 0.75 mg/kg EpoB cohort at 12 weeks post-implantation. Quantitative Imaris 3D volume rendering analysis was performed to determine the volume of myelinated axons. Myelination status was determined by co-localization of MBP with either NF-H+ (total axons) or tdTomato + (CST axons) within the bridge channels **(Fig. 4B, b1-b4)**. We compared this measure between the bridge + Vehicle and Bridge + EpoB combination groups, identifying that EpoB treatment significantly increased the volume of myelinated axons **(Fig. 4C-D)**. These data indicate that EpoB not only promotes axonal growth into the bridge, but that these axons subsequently become myelinated even in the chronic post-injury period, months after the injury and bridge implant.

### DTI analysis reveals improved spinal cord structure with combined bridge and EpoB treatment

Diffusion MRI (dMRI) can be utilized to assess white matter and the orientation of regenerated neural tracts. To determine structural integrity and axonal organization of regenerated tracts within the bridge following Bridge + EpoB treatment, ex vivo DTI was performed in the 1.5 mg/kg EpoB cohort at 12 weeks post-implantation for both bridge groups. Qualitative evaluation of DTI reconstructed spinal cord tractography showed enhanced axonal growth and longitudinal alignment of regenerated tracts within the implantation site in mice receiving the combination Bridge + EpoB group vs. Bridge + Vehicle group; this was apparent in both horizontal **(Fig. 5A and C)** and transverse **(Fig. 5B and D)** projections. For quantitative DTI analysis, we were particularly interested in white matter integrity of descending axons regenerating through the bridge, potentially contributing to locomotor recovery. We therefore used streamline tractography, initiating fiber tract reconstruction (seeding) above the bridge based on anatomical identification of the lesion site. Comparison of Bridge + Vehicle versus Bridge + EpoB treatment groups identified that the combination treatment significantly increased fractional anisotropy (FA) and diffusivity metrics within the lesion/bridge site. Specifically, axial diffusivity (AD) and mean diffusivity (MD) were significantly higher in the Bridge + EpoB group **(Fig. 5E-H)**, a reflection of improved water diffusivity in the tracts. The detection of increased FA and diffusivity in the Bridge + EpoB group likely reflects improved structural integrity of regenerated axons in the chronic SCI lesion.

### CST neurons upregulate growth-state associated proteins following Bridge + EpoB administration

Based on the increased regeneration observed following biomaterial bridge and EpoB administration, we hypothesized that this chronic combinatorial treatment was sufficient to induce CST neurons to adopt a growth-state associated proteomic signature. CST neuronal cell bodies from the ipsilateral and contralateral motor cortex were collected by PALM laser capture microscopy in Crym-tdTomato reporter mice where CST neurons are tdTomato positive **(Fig. 6A)**. Because approximately 90% of CST axons decussate at the level of the pyramids before reaching the spinal cord, we collected layer V tdTomato positive CST cell bodies from the motor cortex in both contralateral and ipsilateral hemispheres. This allows for the evaluation of protein signatures of cells that may project through the injury site versus the intact spinal cord across from the injury site, respectively. Importantly, while this method enabled selective capture of tdTomato positive CST cell bodies, it did not guarantee that all captured cell bodies had initiated growth into the implanted bridge. As a result, this bulk proteome experiment contains data on both regenerated and non-regenerated CST axons, limiting the sensitivity of detection. To maximize sensitivity, all tdTomato-positive CST neuronal cells identified in serial sections through the ipsilateral and contralateral motor cortex, respectively, were captured and pooled for proteomic analysis for each animal analyzed; thus, each animal represented one biological replicate for each of these cortical regions. Using this strategy, we were able to obtain sufficient protein for analysis and identify a regeneration profile for these cells as described below.

Differential expression analysis was performed by comparing EpoB treatment alone vs. Bridge + EpoB treatment to enable separation of the contribution of biomaterial-guided regeneration to the CST proteome and allow EpoB-mediated effects to be distinguished from bridge-driven regenerative changes. This comparison was necessary because EpoB was administered systemically, which would be predicted to induce local sprouting independent of bridge implantation. A total of 2,208 proteins were identified in both groups. As expected, protein signatures were notably distinct across the contralateral and ipsilateral sides of the motor cortex; only 2 proteins displayed conserved downregulation or upregulation across both sides of the cortex **(Fig. 6B-C)**. Volcano plots reveal significantly upregulated (red) and downregulated (blue) proteins on the ipsilateral **(Fig. 6D)** and contralateral hemisphere **(Fig. 6F)**.

These differentially expressed proteins **(Fig. 6D, F)** were compared to a previously published RNA sequencing dataset from Poplawski et al [27] of injured CST neurons that grew into the site of an embryonic stem cell graft and exhibited reactivation of an embryonic transcriptional growth state following SCI. We selected this dataset because the authors collected RNA from CST cells bodies that actively regenerated into the injury site, identified using neuroanatomical tracing, and evaluated transcriptomic signature at 21 DPI, a late regenerative timepoint. Strikingly, 44% (ipsilateral) and 55% (contralateral) of the proteins identified by our mass spectrometry strategy were also identified in the Poplawski et al. 2020 transcriptomic dataset and exhibited parallel directional regulation [27]. **(Fig. 6E, 6G)**. These data highlight the conservation of RNA and protein signatures in growth-state CST neurons and reveal that Bridge + EpoB administration induces an increased regenerative state of CST neurons into the chronic timepoint.

Lastly, we evaluated protein ontology by overrepresentation analysis, which revealed the cellular compartments that were relevant to the differentially expressed proteins. Notably, we found that bridge implantation in combination with EpoB treatment led to increased expression of proteins related to synaptic specialization in the contralateral and ipsilateral motor cortex **(Fig. 6H, 6I)**. Some remodeling of ipsilateral CST motor cortex neurons is expected after a unilateral SCI [28]; consistent with that prediction, ontology analysis of upregulated proteins identified changes related to synaptic membrane and mitochondrial membrane, as well as polysynaptic specialization, and transmembrane transporter complexes **(Fig. 6H)**. Additionally, striking changes were observed in contralateral CST motor cortex neurons, particularly the upregulation of proteins related to postsynaptic density, asymmetric synapses, postsynaptic specialization, and neuron-to-neuron synapses **(Fig. 6I)**. Overall, these data suggest that the combination of bridge and EpoB treatment enhances plasticity in both hemispheres of the motor cortex and drove increased neuronal and synaptic regeneration at the protein level, particularly in contralateral CST neurons.

## Discussion

Regeneration after chronic SCI remains one of the most difficult challenges in neurotrauma, primarily due to the development of dense glial and fibrotic scar tissue and accumulation of growth-inhibitory factors and the progressive decline in intrinsic growth capacity of adult CNS neurons [29-33]. Although biomaterials can support axon growth and locomotor recovery in acute models, their efficacy in chronic settings has been limited [12]. Our prior work showed that acute implantation of porous, biodegradable multichannel bridges promoted robust axonal regeneration, including regeneration of the highly refractory CST, re-establishment of cortical-motor relay circuits, and significant functional recovery [5, 6]. However, similar bridge implantation in chronic injury failed to elicit meaningful functional improvement [12], highlighting the need to address both the intrinsic growth potential of injured neurons and the extrinsic inhibitory microenvironment.

Microtubule stabilization with drugs such as Taxol and epothilone drugs (EpoB and EpoD) has been shown to reduce fibrotic and glial scarring, decrease accumulation of extracellular matrix inhibitors, and reinitiate a growth-competent state in injured axons, resulting in improved motor outcomes after SCI [15, 16, 19, 30, 34, 35]. Of note, the molecular mechanism of epothilone B induced axon regeneration has been recently described [36]. Indeed, a recent study with chronic PLG bridge implantation in combination with Epothilone D (EpoD) administration significantly lowered fibronectin and GFAP scar tissue formation [19]. However, a critical gap in the field is to promote regeneration of long tracts, particularly the CST, a tract that regenerates poorly [37-39], in the chronic post-injury period. Accordingly, in this study, we investigated whether combining a biomaterial bridge to guide axons that have reinitiated growth with systemic EpoB administration would enable CST regeneration and functional recovery in a chronic SCI paradigm.

We report that, in two independent chronic SCI cohorts (EpoB = 0.75 mg/kg and 1.5 mg/kg), bridge implantation combined with EpoB treatment induced significant, dose-dependent improvement in locomotor recovery. While both EpoB doses yielded improvement when combined with bridges, 1.5 mg/kg exhibited a trend towards a more pronounced recovery. These data suggest that the 1.5 mg/kg dose provided more robust and sustained microtubule stabilization, thereby accelerating axonal extension and synaptic reconnection. While the bridge alone supported modest axonal ingrowth, the addition of EpoB significantly increased both total (NF-H^+^) and CST (Crym-tdTomato^+^) axonal volume, and the volume of myelinated axons, in the bridge. Although previous in vitro studies have reported that EpoB can inhibit oligodendrocyte precursor cell differentiation and may induce dysmyelination in neuron-oligodendrocyte co-cultures [40], our in vivo data demonstrate that the capacity for myelination is retained at the EpoB doses used in this study. This outcome likely reflects the recruitment of both oligodendrocytes and PNS-derived Schwann cells, both of which infiltrate SCI lesions [41], which would be consistent with the improved diffusivity parameters detected by DTI and enhanced axonal integrity and tract coherence. Whether the addition of strategies specific to driving myelination could further drive recovery is a subject for additional investigation.

DTI MRI is a clinical imaging modality used in assessing spinal cord integrity [42]. In preclinical studies, DTI provides a whole-tissue analysis of white matter integrity and axonal organization that can complement and validate histological findings. While in vivo DTI would allow longitudinal monitoring of lesion evolution and treatment effects [43], ex vivo imaging affords superior signal-to-noise ratio, eliminates motion artifacts, and supports high-resolution, extended acquisitions [44]. Despite inherent analytical challenges associated with fixed-tissue DTI, such as reduced diffusivity values and potential partial-volume effects, our analysis revealed that the Bridge + EpoB combination group exhibited significantly higher fractional anisotropy, axial, and mean diffusivity within the lesion/bridge region, which are broad indicators of axon and tract integrity. These findings align with histological evidence of increased regeneration and remyelination, suggesting that the combination of Bridge + EpoB promotes structurally integrated repair in chronic SCI.

Lastly, we investigated the protein signatures of CST cell bodies using laser capture microdissection of Crym-tdTomato positive neurons in the motor cortex, followed by unbiased mass spectrometry. As the longest tract in the nervous system, CST neurons originate in the motor cortex [45], decussate in the caudal portion of the brain, and innervate the contralateral muscles to control movement. Approximately 90% of CST axons decussate to the contralateral spinal cord, whereas 10% are retained to project ipsilaterally [28]. Consistent with this projection ratio, we identified changes in the proteome of both ipsilateral and contralateral CST neurons. Bridge + EpoB treatment notably reactivated molecular programs associated with active growth states and increased synaptic proteins in contralateral motor cortex neurons. We also identified increased regeneration and synaptic proteins in the cortical neurons of the ipsilateral hemisphere, an observation that could reflect systemic administration of EpoB in this study. Most interestingly, proteins identified as upregulated in contralateral motor cortex neurons in the Bridge + EpoB combination vs. EpoB alone group exhibited a striking parallel with transcriptomic data for pro-regenerative CST neurons [27].

Although few transcriptomic studies have defined gene expression programs associated with CST injury and regeneration [27, 39, 46-50], we particularly leveraged the dataset from Poplawski et al., which investigated regeneration-associated gene expression in CST neurons that extend axons into a transplanted cell graft at 21 DPI and demonstrated that CST regeneration is accompanied by a reversion to an embryonic-like transcriptional state. These two datasets represent completely different avenues of evidence suggesting activation of a regenerative program within these cells. Strikingly, our proteomic analysis revealed substantial overlap with these regeneration-associated transcriptomic signatures, despite inducing CST regeneration at a much later time point (60 DPI) using our combinatorial Bridge + EpoB therapy. This correspondence between RNA-level and protein-level signatures suggests that adult CST neurons retain the ability to re-engage developmental growth programs when provided with appropriate extrinsic cues.

Together, our findings demonstrate that combining structural guidance from bridges with EpoB-mediated neuronal reactivation enables meaningful CST regeneration and functional recovery even in the chronically injured spinal cord, offering a promising therapeutic avenue for patients with established SCI.

## AUTHOR CONTRIBUTIONS

<u>Conceptualization and study design:</u> A.J.A., A.A., U.N., B.J.C., L.D.S., and F.B.; Methodology: A.J.A., U.N., A.A., P.S.S., R.A.N., C.M.D., S.M., H.S., A.O., J.B.L., P.D.G., and F.B.; <u>Investigation and analysis:</u> In vivo animal experiments by A.A., U.N., and R.A.N.; <u>Immunohistochemistry, imaging, and quantification:</u> by U.N., and S.M.; <u>DTI acquisition and analysis</u>: A.O., and J.B.L.; <u>PALM laser capture and microdissection</u>: R.A.N., P.S.S., A.A., and H.S.; <u>Mass spectrometry experiments and analysis</u>: P.D.G. and P.S.S.; <u>Biomaterial bridge fabrication</u> by C.M.D. and L.D.S.; <u>Epothilone B dosing optimization</u> by F.B. <u>Manuscript writing and figure assembly</u>: U.N., <u>Manuscript review and editing</u>: P.S.S., and A.J.A.; <u>Supervision:</u> A.J.A., B.J.C., and L.D.S.; <u>Funding acquisition:</u> A.J.A., B.J.C., L.D.S., and P.D.G.

## Acknowledgments

We thank Javier Lepe and Chris Nelson for assistance with animal surgeries. We gratefully acknowledge George Mina for support with behavioral data analysis and Rohan Alexander Webb for assistance with histological analysis. This work was supported by the National Institutes of Health R01NS117103 and R01EB005678. F.B. is supported by the International Foundation for Research in Paraplegia, Wings for Life, ERANET AxonRepair, ERANET RATER SCI and Chan-Zuckerberg Initiative (CZI). F.B. is also funded by the Deutsche Forschungsgemeinschaft (DFG, German Research Foundation) - Project-ID 227953431 – SFB 1089, as well as SFB 1158, SFB 1690, and SPP 2395. F.B. is a member of the excellence cluster ImmunoSensation2 (EXC2151– 390873048) and the iBehave NRW network. F.B. is a recipient of the Roger de Spoelberch Prize. Proteomic analysis of corticospinal neurons by mass spectrometry was supported by NIH instrumentation grant S10 OD016328 to P.D.G. Figure 1 was created in BioRender https://BioRender.com/le0nbhp

## Data availability

Data will be made available on request.

